# Metapopulation ecology links antibiotic resistance, consumption and patient transfers in a network of hospital wards

**DOI:** 10.1101/771790

**Authors:** Julie Teresa Shapiro, Gilles Leboucher, Anne-Florence Myard-Dury, Pascale Girardo, Anatole Luzatti, Mélissa Mary, Jean-François Sauzon, Bénédicte Lafay, Olivier Dauwalder, Frédéric Laurent, Gérard Lina, Christian Chidiac, Sandrine Couray-Targe, François Vandenesch, Jean-Pierre Flandrois, Jean-Philippe Rasigade

## Abstract

Antimicrobial resistance (AMR) is a global threat. A better understanding of how antibiotic use and between ward patient transfers (or connectivity) impact hospital AMR can help optimize antibiotic stewardship and infection control strategies. Here, we used metapopulation ecology to explain variations in infection incidences of 17 ESKAPE pathogen variants in a network of 357 hospital wards. Multivariate models identified the strongest influence of ward-level antibiotic use on more resistant variants, and of connectivity on nosocomial species and carbapenem-resistant variants. Pairwise associations between infection incidence and the consumption of specific antibiotics were significantly stronger when such associations represented a priori AMR selection, suggesting that AMR evolves within the network. Piperacillin-tazobactam consumption was the strongest predictor of the cumulative incidence of infections resistant to empirical sepsis therapy. Our data establish that both antibiotic use and connectivity measurably influence hospital AMR and provide a ranking of key antibiotics by their impact on AMR.

---

Antimicrobial resistance (AMR) of pathogenic bacteria progresses worldwide and imposes a considerable burden of morbidity, mortality and healthcare costs^1,2^. AMR is increasingly recognized to emerge in various settings including agriculture^3^ or polluted environments^4,5^. However, hospitals remain major hotspots of AMR selection^6,7^ that concentrate strong antibiotic pressure, fragile patients and highly resistant pathogens^8^, while patient transfers between wards and facilities accelerate pathogen dissemination^9^.

The primary hospital-based strategies against AMR are antimicrobial stewardship and infection control^10^, which aim respectively to lower the antibiotic pressure and the transmission of pathogens. The need for such strategies is widely accepted^11^ but their implementation details are more debated^12^, especially regarding which antibiotics should be restricted first^13^ or the risk-benefit balance of screening-based patient isolation procedures^14,15^. Thus far, designing efficient antibiotic stewardship strategies has been hindered by the paucity of evidence concerning which antibiotics exert the strongest selection pressure. As a result, available rankings of antibiotics for de-escalation and sparing strategies rely on expert consensus with partial agreement^16^, themselves based on conflicting evidence^17,18^.

Linking antibiotic use and AMR prevalence is difficult due to the confounding effects of bacterial transmission and the complexity of the ecological processes underlying AMR (reviewed in^19^). Observational studies of AMR usually report on the proportion of resistant variants in a limited set of species, which conceals the overall burden of AMR and can make interpretation difficult when, for instance, resistant variants apparently increase in proportion while decreasing in incidence^20^. To alleviate these issues, studies of the impact of antibiotic use on AMR in hospitals could benefit from ecological frameworks able to simultaneously model the incidence of infection with most relevant pathogens while controlling for confounding effects. Metapopulation ecology is such a framework. It was introduced by Levins^21^ to explain the persistence of agricultural pests across a set of habitat patches and refined by Hanski to account for the size of patches, the connectivity between them, and habitat quality within them^22,23^. Metapopulation models, beyond their frequent use in wildlife and conservation biology^24–26^, have recently provided theoretical grounds for pathogen persistence in the healthcare setting^27^. So far, however, metapopulation models of hospital AMR have been applied on simulated rather than empirical data^27,28^.

We used metapopulation modeling to isolate the effect of antibiotic consumption on the incidence of infections with 7 major pathogen species and their resistant variants within a 5,400-bed, 357-ward hospital network, using detailed data over the course of one year. We considered that bacteria in this network approximate a metapopulation of local populations in patches represented by hospital wards, connected by transfers of colonized patients^29^.

Our primary objective was to determine the respective impacts of antibiotic use and connectivity on the incidence of infections with resistant pathogens. The secondary objective was to compare the impacts of specific antibiotics on hospital AMR, by modeling each pathogen variant incidence separately, then by considering all variants resistant to drugs commonly used in empirical sepsis therapy. Our findings highlighted both common patterns and species-specific behaviors of pathogens and identified a major, so far underappreciated, association between the widely-used drug piperacillin-tazobactam and resistance to both 3^rd^-generation cephalosporins and carbapenems.

## Results

### Distribution of bacterial pathogens and antibiotic use in a hospital network

We analyzed pathogen isolation incidence in clinical samples, antibiotic use and patient transfers in 357 hospital wards from the region of Lyon, France, from October 2016 to September 2017. Ward-level data were aggregated from 14,034 infection episodes, defined as ward admissions with ≥1 clinical sample positive for *E. coli* or an ESKAPE pathogen (*Enterococcus faecium*, *Staphylococcus aureus*, *Klebsiella pneumoniae*, *Acinetobacter baumannii*, *Pseudomonas aeruginosa* and *Enterobacter cloacae* complex), collectively termed ESKAPE_2_. Pathogens were grouped into species-resistance pattern combinations, namely 3^rd^-generation cephalosporin-resistant E. coli, E. cloacae complex and K. pneumoniae (3GCREC, -EB and -KP), carbapenem-resistant *E. coli*, *E. cloacae* complex, *K. pneumoniae*, *P. aeruginosa* and *A. baumannii* (CREC,-EB, -KP, -PA and -AB), vancomycin-resistant *E. faecium* (VREF) and methicillin-resistant S. aureus (MRSA). Pathogen variants not falling into these resistance groups were designated by species (EC, EB, KP, PA, AB, EF, SA) and collectively referred to as the less-resistant variants.

Infection episodes most frequently involved the less-resistant variants, mainly EC, SA, PA and KP (Table 1). These variants were also found in the largest number of wards. Resistant variants were consistently less frequent than their less-resistant variants in all species and, in enterobacteria, carbapenem resistant variants were consistently less frequent than 3GC-resistant variants. VREF and CRAB infections were exceptional.

**Table 1.** Distribution of ESKAPE_2_ pathogen infection episodes in 357 hospital wards.

| Taxon | Resistance profile | Acronym | No. of episodes<br>(%), n=14,034 | No. of wards<br>(%), n=357 | Concentration index<sup>a</sup> (%) (95% CI) |
| --- | --- | --- | --- | --- | --- |
| *Escherichia coli* | 3GC- and carbapenem-S | EC | 6,308 (44.9) | 328 (91.9) | 0.6 (0.6 – 0.7) |
|  | 3GC-R | 3GCREC | 736 (5.2) | 207 (58.0) | 0.7 (0.6 – 0.8) |
|  | Carbapenem-R | CREC | 24 (0.2) | 24 (5.6) | 1.4 (0.0 – 3.5) |
| *Klebsiella pneumoniae* | 3GC- and carbapenem-S | KP | 1,134 (8.1) | 249 (69.7) | 0.7 (0.6 – 0.8) |
|  | 3GC-R | 3GCRKP | 530 (3.8) | 175 (49.0) | 0.9 (0.7 – 1.0) |
|  | Carbapenem-R | CRKP | 43 (0.3) | 32 (9.0) | 1.7 (0.0 – 3.8) |
| *Enterobacter cloacae* complex | 3GC- and carbapenem-S | EB | 279 (2.0) | 140 (39.2) | 1.0 (0.7 – 1.3) |
|  | 3GC-R | 3GCREB | 212 (1.5) | 116 (32.5) | 0.8 (0.5 – 1.0) |
|  | Carbapenem-R | CREB | 102 (0.7) | 74 (20.7) | 0.7 (0.3 – 1.0) |
| *Pseudomonas aeruginosa* | Carbapenem-S | PA | 1,188 (8.5) | 231 (64.7) | 0.8 (0.7 – 0.9) |
|  | Carbapenem-R | CRPA | 444 (3.2) | 148 (41.5) | 1.5 (1.2 – 1.7) |
| *Acinetobacter baumannii* | Carbapenem-S | AB | 96 (0.7) | 61 (17.1) | 1.3 (0.6 – 2.1) |
|  | Carbapenem-R | CRAB | 12 (0.1) | 10 (2.8) | 3.0 (0.0 – 9.4) |
| *Enterococcus faecium* | Vancomycin-S | EF | 503 (3.6) | 133 (27.3) | 1.4 (1.2 – 1.6) |
|  | Vancomycin-R | VREF | 7 (<0.1) | 7 (2.0) | 0.0 (0.0 – 9.5) |
| *Staphylococcus aureus* | Methicillin-S | SA | 2,113 (15.1) | 273 (76.5) | 0.7 (0.5 – 0.9) |
|  | Methicillin-R | MRSA | 303 (2.2) | 151 (42.3) | 0.7 (0.6 – 0.9) |

NOTE. <sup>a</sup>The concentration index estimates the probability that two episodes taken at random occurred in the same ward. 3GC, 3<sup>rd</sup>-generation cephalosporins; S, susceptible; R, resistant.

To estimate the degree of concentration of each variant in the network, we calculated concentration indices defined as the probability that two random occurrences of the same variant originated from the same ward, analogous to the asymptotic Simpson index (see Methods). The concentration of infection episodes was weak (<5%) for all variants, indicating a global lack of clustering (Table 1). Concentration increased with resistance (~2-fold increase from the least to the most resistant variant) in *E. coli*, *K. pneumoniae*, *P. aeruginosa* and *A. baumannii*, suggesting an adaptation of resistant variants to more specific habitats compared to their less-resistant counterparts. This pattern was not found in *E. cloacae* complex, *E faecium* and *S. aureus*.

Antibiotics were prescribed in 86.3% of wards (Table 2), with a total consumption of 125.7 defined daily doses per year per bed (ddd/y/b). Antibiotics usually suspected to select for AMR in the selected ESKAPE_2_ variants were grouped into 9 classes (Table 2). Antibiotics with comparatively rare use (e.g., rifampicin), narrow spectrum (e.g. amoxicillin) or most frequently used in combination therapy (aminoglycosides) were excluded. The distribution of antibiotic use in the network was analysed using the concentration index described above, here representing the probability that two random doses were delivered in the same ward. Antibiotic use was diffuse, with concentration indices <4%, ranging from 0.8% for CTX/CRO and FQ to 3.6 for OXA.

**Table 2.**
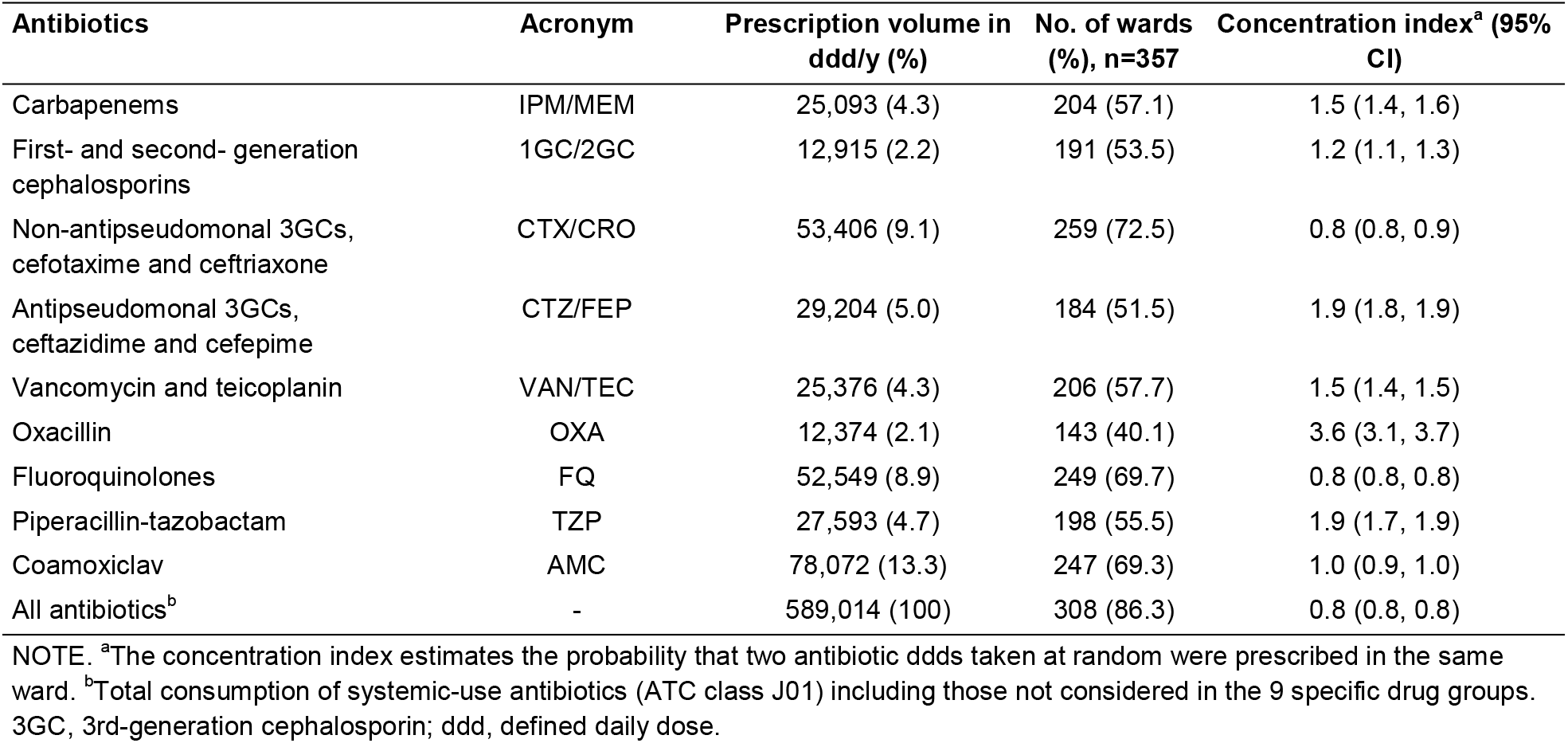
Distribution of antibiotic use in hospital wards.

### Antibiotic use and connectivity predict ESKAPE_2_ infection incidence

We used generalized linear models (GLMs) within the metapopulation framework to disentangle the influences of antibiotic pressure, connectivity and other ward characteristics on the incidence of infections with ESKAPE_2_ pathogens and their resistant variants. Connectivity quantifies the incoming flux of each pathogen variant in a downstream ward receiving infected patients from upstream wards. Practically, we estimated connectivity for each variant and downstream ward as the sum of the transfers from each upstream ward multiplied by the variant’s prevalence in that ward (see Methods). Wards were characterized by their size (no. of beds) and type, representing patient fragility and coded on an ordinal scale with a score of 2 for intensive care and blood cancer units, 1 for progressive care units and 0 for other wards. Importantly, the observed incidence in a ward depends directly on the frequency of sampling and specimen types (e.g., respiratory vs urinary tract specimens), leading to sampling bias. To correct for this bias, all models included a baseline incidence value as a control covariate, defined as the ward-level incidence predicted by sampling alone, assuming that the probability of pathogen presence in a given sample is constant across wards (see Methods). As expected, the control value was strongly correlated with both antibiotic use and the incidence of infections in all prevalent variants (Supplementary Fig 1 and 2). The incidence of each pathogen variant was then modelled as a separate Poisson-distributed GLM (Figure 1) adjusted for sampling bias. [In unadjusted, bivariate analysis, the incidence of each variant was strongly correlated with both antibiotic use (Supplementary Figure 3) and connectivity (Supplementary Figure 4), with the exception of the very rare CRAB and VREF variants.]

**Figure 1.**
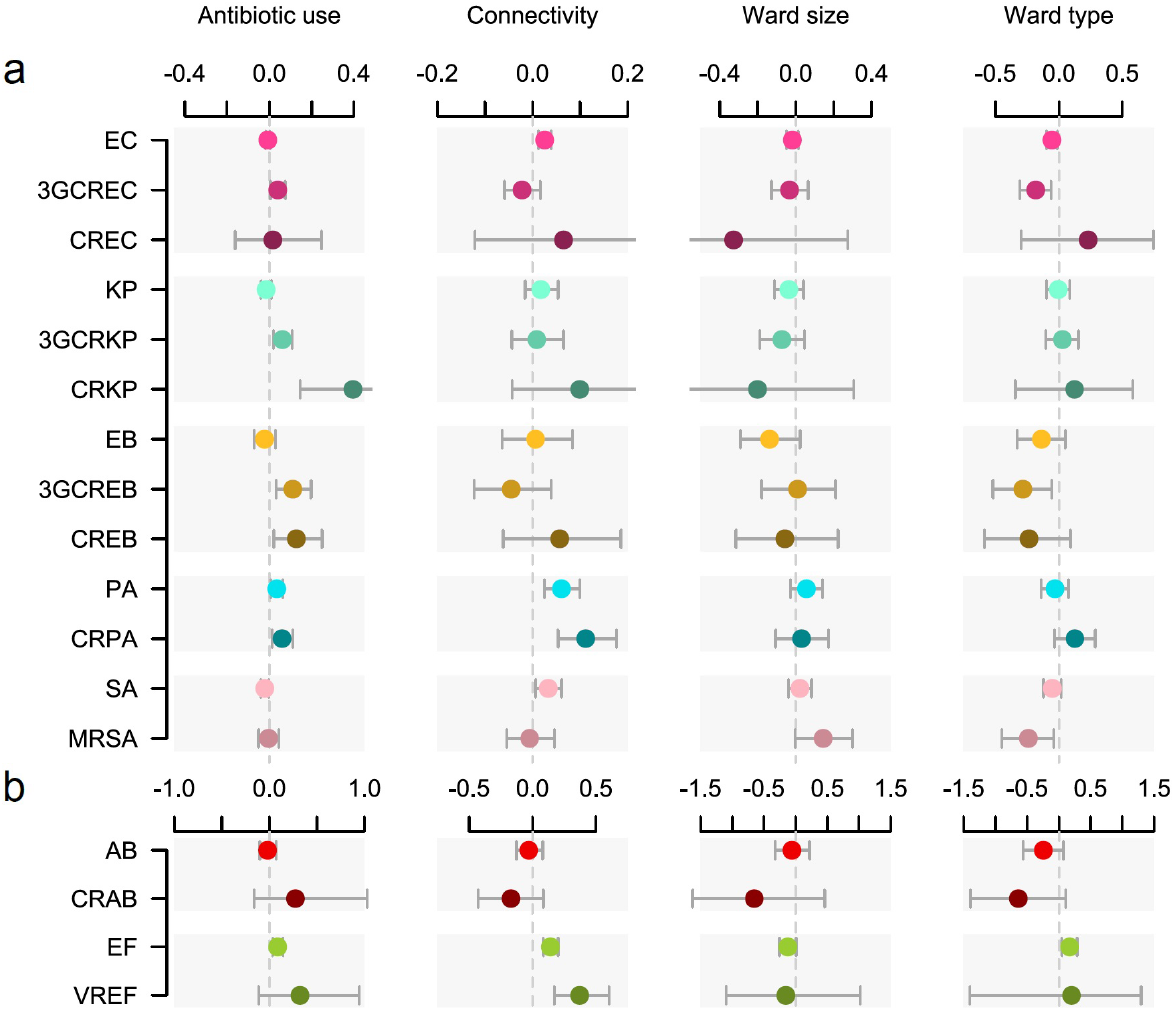
Antibiotic use and connectivity predict the incidence of infection with ESKAPE2 pathogen variants. Shown are the 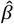 coefficients (points) and 95% confidence intervals (bars) of Poisson regression models of the incidence of each variant in each ward (n=357) for antibiotic use, connectivity (estimated no. of patients colonized with the same variant entering the ward), ward size (no. of beds) and ward type, coded 2 for intensive care and blood cancer units, 1 for progressive care units and 0 for other wards. All models were adjusted for sampling bias. Models involving A. baumannii and E. faecium, which exhibited larger 95% CIs due to smaller incidence of the resistant variants, are shown with separate scales (panel b) for readability. Coefficients approximately represent the relative change of incidence of infections per doubling of the predictor.

In the adjusted GLMs, global antibiotic use was significantly associated with infection incidence in 8 pathogen variants independent of connectivity, ward size and type (Figure 1). The maximum effect size was found in CRKP with a coefficient of 0.4, which predicted approximately a 50% increase of incidence per 2-fold increase of antibiotic use. Strikingly, the magnitude of association of antibiotic use with incidence increased with resistance levels within all taxa excepted CREC (Figure 1). This pattern supported a general influence of antibiotic use on within-species AMR. Antibiotic use was significantly associated with a decreased incidence in only one variant, namely SA.

The influence of connectivity on infection incidence was more variant-specific than that of antibiotic use. Connectivity had a positive effect for two resistant variants (CRPA and VREF) and four less-resistant variants (EC, PA, SA, EF). The effect unambiguously increased with resistance in *P. aeruginosa* and *E. faecium* but not in other species, although connectivity exhibited comparatively larger effects for carbapenem-resistant variants in enterobacteria (Figure 1). The effect of ward size was weaker and generally insignificant compared to antibiotic use and connectivity. Notably, ward type had a negative, significant effect in four variants (EC, 3GCREC, 3GCREB, and MRSA), suggesting that these infections preferentially occur in general wards with less fragile patients.

### Possibly causal associations between antibiotic use and resistance

The metapopulation models illustrated in Figure 1 identified positive associations between total antibiotic use in hospital wards and increased incidences of infections with the more resistant variants of several ESKAPE_2_ species. Yet, a correlation with AMR does not establish a causal role of antibiotics. For instance, a high incidence of resistant infections in a ward can increase antibiotic use through prolonged or combined therapies^19^. Conversely, the prescription of antibiotics always inactive against a variant is unlikely to be motivated by this variant’s incidence and such antibiotics are more likely to provide a direct benefit to the resistant variant. Based on this rationale, we propose stringent criteria to identify possibly causal associations between specific antibiotics and pathogen variants (see Methods). Under the hypothesis that antibiotic use is either a consequence of AMR or spuriously correlated with AMR, the strength of an association between the use of an antibiotic and the incidence of a variant should not depend on whether the association fulfills the criteria for possible causality.

To test this hypothesis, we identified possibly causal associations in our data and examined whether they were equally likely to be positive and significant compared to other associations. We constructed Poisson regression models where the total antibiotic use was replaced with the use of specific antibiotics, along with the incidence control and connectivity covariates (Figure 2a). These 17 variant-specific models, each built using 9 antibiotic groups as predictors, yielded 153 coefficients of which 17 (11.1%) represented possibly causal associations (see Methods). Positive and significant coefficients were found in 6/17 possibly causal associations (35.3%), more frequently so than in other associations (10/136, 7.4%) with an odds-ratio of 6.7 (95% CI, 1.7 to 25.6; p=0.003, Fisher’s exact test, two-sided). The median regression coefficient in possibly causal associations was also significantly higher than in other associations (p=0.009, Mann-Whitney *U*-test, two-sided; Figure 2b). Four of the 6 significant, possibly causal associations involved CTX/CRO, which selected for 3GCREC, 3GCRKP, PA and EF; the other two involved carbapenems selecting for CRPA, and AMC selecting for MRSA. Overall, the enrichment of possibly causal associations among significant model coefficients confirmed that the local, ward-level selection of drug-resistant variants by antibiotics is measurably pervasive throughout our hospital network.

**Figure 2.**
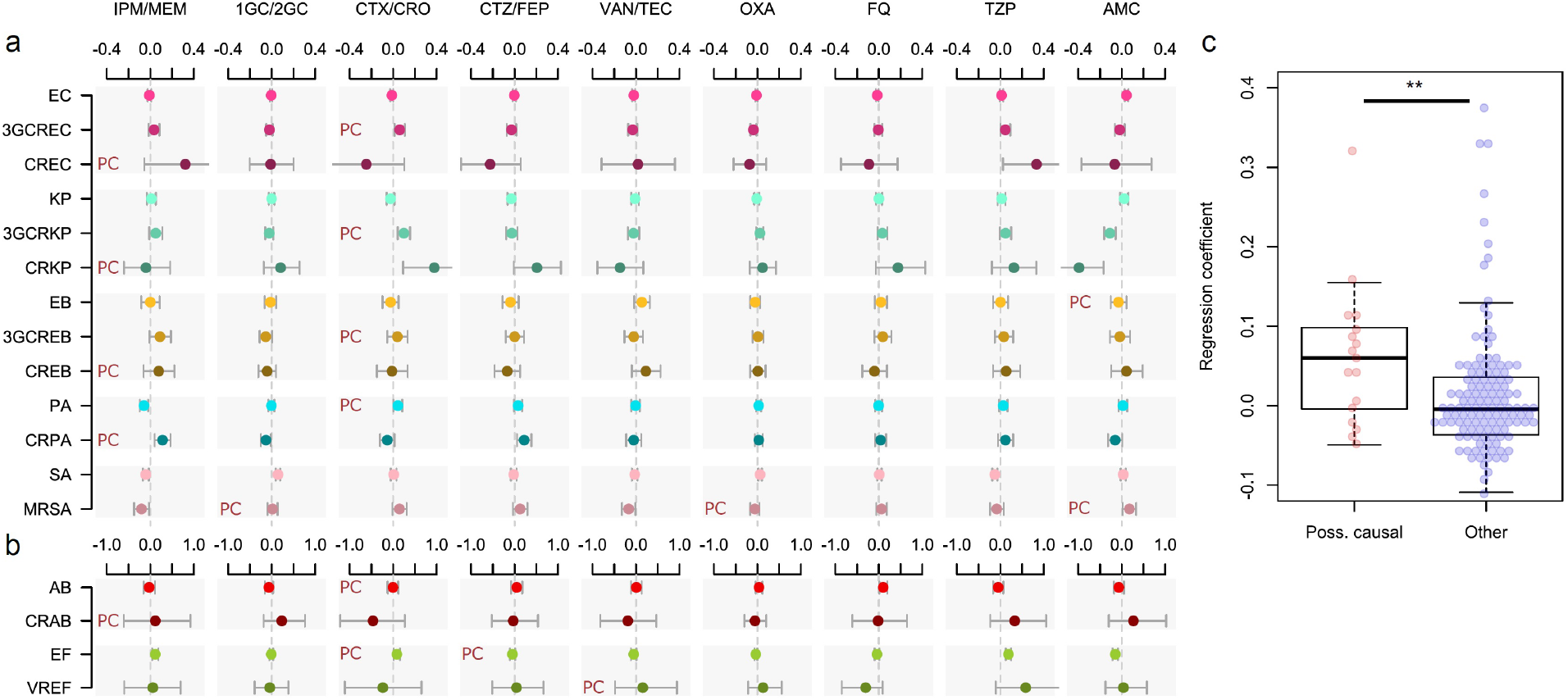
Possibly causal associations between the use of specific antibiotics and the incidence of infection with ESKAPE2 pathogen variants. Shown are the 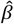 coefficients (points) and 95% confidence intervals (bars) of Poisson regression models of the incidence of each variant in each ward (n=357) for consumption volumes of 9 antibiotic groups, after controlling for sampling bias and connectivity. Associations classified as possibly causal (n=17) are indicated by a ‘PC’ mark. Models involving *A. baumannii* and *E. faecium*, which exhibited larger 95%CIs due to smaller incidence of the resistant variants, are shown with separate scales (panel b) for readability. (c), the coefficients of possibly causal associations are significantly higher (p<0.01, Mann-Whitney *U*-test, two-sided) than the coefficients of other associations. The center line indicates the median; box limits indicate the upper and lower quartiles; whiskers indicate the 1.5× interquartile range; points indicate the individual coefficients.

### Quantifying the drivers of cefotaxime/ceftriaxone and carbapenem resistance

From a clinical stand-point, the most immediate consequence of AMR is the failure to control sepsis with empirical antibiotics, mainly carbapenems and non-antipseudomonal 3GCs such as cefotaxime. Because such failure can equally result from acquired or intrinsic resistance, the incidence of intrinsically resistant pathogens such as EF is of equal clinical importance as that of pathogen variants with acquired resistance mechanisms. To examine the impact of antibiotics on both intrinsic and acquired resistance, we modeled the cumulative incidence of infections with 3GC- and/or carbapenem-resistant (3GCR and CR) variants of the ESKAPE_2_ pathogens (see Methods).

In these models, antibiotic use was not only the sole predictor with a positive effect on both CR and 3GCR incidence, but also the strongest predictor (Figure 3a). Connectivity predicted CR, but not 3GCR, incidence, in line with the comparatively greater impact of connectivity on individual CR variants (Figure 1). Ward size had no measurable effect in either model. Ward type, reflecting patient fragility, had no effect on CR incidence but was negatively associated with 3GCR incidence, in line with similar associations found for 3GCREC and 3GCREB (Figure 1). These findings provide an unambiguous link between antibiotic use and global resistance to empirical sepsis therapy that was robust to confusion by sampling bias, connectivity and other ward characteristics.

**Figure 3.**
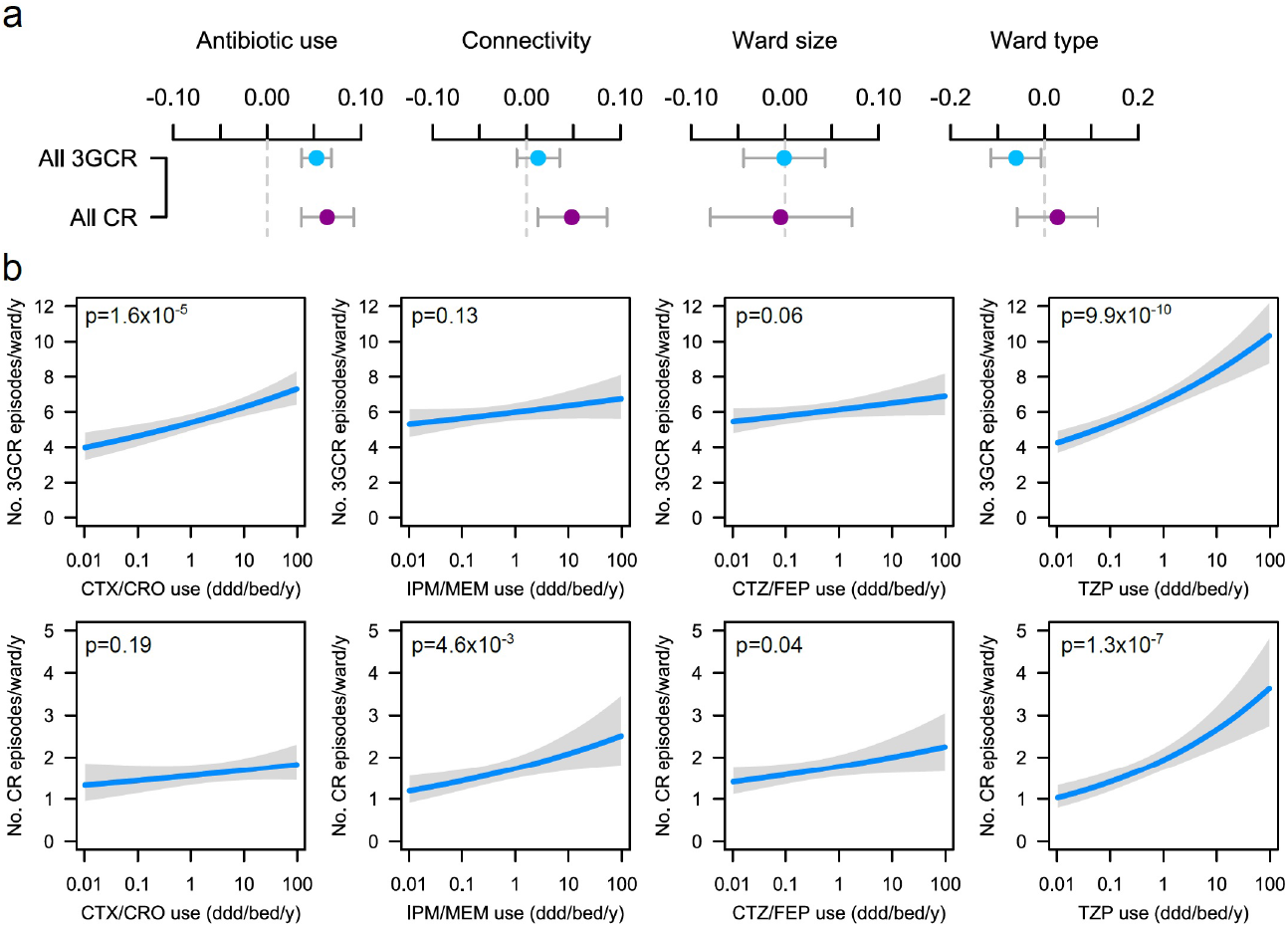
Global and specific antibiotics consumption predict the incidence of infection with 3^rd^-generation cepalosporin- or carbapenem-resistant ESKAPE2 variants. (a), 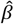 coefficients (points) and 95% confidence intervals (bars) of Poisson regression models of the incidence of 3GCR and CR infections in each ward (n=357). Models were adjusted for sampling bias. (b) Predicted incidence and 95% confidence bands of infections with 3GCR and CR infections, as a function of CTX/CRO, IPM/MEM, CTZ/FEP and TZP consumption in models adjusted for sampling bias, connectivity and the consumption of 5 other antibiotic groups. Variants classified as 3GCR were 3GCREC, 3GCRKP, CRKP, 3GCREB, CREB, PA, CRPA, AB, CRAB, EF, VREF and MRSA; the CR category included CREC, CRKP, CREB, CRPA, CRAB, EF, VREF and MRSA.

To determine which antibiotics have the strongest impact on global carbapenem and 3GC resistance, we examined the effect of replacing the total antibiotic use in our models by individual antibiotics, similar to the approach described in the previous section. Antibiotics whose use significantly predicted either 3GCR or CR incidence were CTX/CRO, IPM/MEM, CTZ/FEP and TZP (Table 3). To visualize their respective impact, we plotted the average ward-level infection incidence predicted by variations of the consumption volumes in the models adjusted for connectivity and sampling bias (Figure 3b). The resulting pattern of association highlighted, again, possibly causal associations: 3GCR incidence was significantly predicted by the consumption of CTX/CRO but not IPM/MEM, while CR incidence was significantly predicted by the consumption of IPM/MEM, but not CTX/CRO. CTZ/FEP had a barely significant influence on CR incidence. Strikingly, TZP consumption predicted both 3GCR and CR infection incidences and, in both models, TZP coefficients outweighed other coefficients by a large margin in terms of amplitude and significance. Overall, these results demonstrate a specific effect of CTX/CRO and IPM/MEM consumption on resistance to the same antibiotic group, but not other groups, and identify a major role of TZP consumption in predicting the incidence of both 3GCR and CR infections. To propose a unified ranking of the impact of antibiotics on 3GC and CR resistance, a final model was constructed by pooling all 3GCR and CR variants together (Table 3). In this model, TZP and CTX/CRO had positive and significant coefficients; CTZ/FEP had positive but insignificant coefficients; FQ and OXA had negative, insignificant coefficients; and VAN/TEC, 1GC/2GC and AMC had negative and significant coefficients.

**Table 3.** Coefficients of regression of the incidence of 3GCR and/or CR infections for the consumption of 9 antibiotic groups in 357 wards.

| Antibiotics | Regression coefficient<sup>a</sup> x100 (95% CI) |  |  |
| --- | --- | --- | --- |
|  | 3GCR incidence model | CR incidence model | 3GCR or CR incidence model |
| TZP | 6.30 (4.28, 8.32) | 9.56 (6.02, 13.12) | 6.42 (4.41, 8.44) |
| CTX/CRO | 4.59 (2.50, 6.66) | 2.43 (-1.24, 6.09) | 4.47 (2.39, 6.54) |
| CTZ/FEP | 1.89 (-0.05, 3.83) | 3.54 (0.09, 7.02) | 1.78 (-0.15, 3.73) |
| IPM/MEM | 1.67 (-0.51, 3.85) | 5.62 (1.75, 9.54) | 1.78 (-0.40, 3.96) |
| FQ | -0.19 (-1.86, 1.48) | -1.39 (-4.31, 1.53) | -0.21 (-1.87, 1.46) |
| OXA | -0.42 (-1.59, 0.75) | -1.77 (-3.71, 0.18) | -0.49 (-1.65, 0.68) |
| VAN/TEC | -2.26 (-4.20, -0.30) | -3.63 (-7.04, -0.20) | -2.21 (-4.15, -0.26) |
| 1GC/2GC | -2.42 (-3.78, -1.07) | -2.80 (-5.14, -0.44) | -2.44 (-3.78, -1.09) |
| AMC | -4.17 (-6.23, -2.09) | -5.51 (-9.06, -1.91) | -4.18 (-6.24, -2.11) |

NOTE. <sup>a</sup>Coefficients were obtained from Poisson regression models adjusted for sampling bias, connectivity and ward size and type.

## Discussion

The impact of antibiotic use on resistance has been demonstrated at different scales including hospitals, regions and countries^30–32^, but a quantitative assessment of this impact within hospitals is still lacking. By applying metapopulation ecology to explain variations of ESKAPE_2_ infection incidences across a large network of hospital wards, we demonstrate that both antibiotic use and inter-ward patient transfers independently contribute to ward-level AMR in several species. Our study also provides the first quantitative ranking of the impact of several key antibiotics on the global burden of drug-resistant ESKAPE_2_ infections in a hospital network.

Previous theoretical work based on modeling and simulation has predicted how patient transfers contribute to AMR prevalence through pathogen dissemination^33–35^. Our study provides an empirical confirmation of these predictions and identifies the species and variants most influenced by connectivity (Figure 1). Understanding the respective impacts of antibiotic use and connectivity on the burden of resistant infections is essential for optimizing interventions against AMR. If the resistant infections in a ward mostly result from the admission of already colonized patients, one would expect antibiotic restrictions to have a limited impact on AMR compared to infection control measures to prevent the further dissemination of the pathogens. Conversely, a weak influence of connectivity suggests that resistant pathogens are either selected locally or introduced from sources outside the network. Consequently, connectivity’s influence on incidence should be higher in pathogen variants endemic in the hospital, and lower in community-associated variants or variants whose resistance is selected locally. This model is consistent with our finding that connectivity had the strongest influence in the typical nosocomial pathogens *P. aeruginosa* and *E. faecium* and a comparatively lower influence in community-associated variants (3GCREC and −KP). The low influence of connectivity on the incidence of resistant *E. cloacae* complex variants can be explained by their local selection. While resistance in *E. coli* and *K. pneumoniae* typically requires gene acquisition^34,35^, *E. cloacae* complex can resist cephalosporins and carbapenems through increased AmpC betalactamase and decreased porin expression^36–38^. Such resistance emerges through adaptation and *de novo* mutations that are rapidly selected from the local reservoir of susceptible progenitors^39,40^. In contrast, the incidence of global CR infections was strongly influenced by connectivity (Figure 3), with an effect size comparable to that of antibiotic consumption, while connectivity had no measurable influence on global 3GCR infections. This suggests that infection control measures could be particularly effective at preventing the spread of CR pathogens between wards.

The rationale behind hospital-based antibiotic stewardship is based on the assumption that AMR evolves in hospitals. Yet, there is surprisingly limited evidence to support this assumption^36–38^. Ecological studies have repeatedly identified associations between the use of antibiotics and AMR prevalence, but such associations do not necessarily reflect AMR selection^19^. However, more specific associations at the antibiotic and variant level are not all equally likely to be causal or consequential. Based on medical and biological reasoning, we identified associations representing possible selection and showed they outweighed other associations in terms of significance and amplitude (Figure 2). Other significant associations could not be classified as possibly causal but might reflect co-selection. The association of CTX/CRO with CRKP but not CREC likely reflected, in our setting, the selection for the highly frequent 3GC-resistance in carbapenemase-producing K. pneumoniae, contrasting with *E. coli* Oxa_48_ producers that frequently remain 3GC-susceptible but TZP-resistant^39^, consistent with the strong association between TZP use and CREC incidence. These findings confirm that the associations between antibiotic use and the incidence of specific resistant variants are preferentially causal in nature and, consequently, that AMR evolves in our hospital network.

Because intrinsic and acquired resistances to an antibiotic equally lead to treatment failure, we modeled the pooled incidences of infections with 3GC- or carbapenem-resistant variants of the ESKAPE_2_ pathogens, including those with intrinsic resistance. This approach allowed us to rank antibiotics by their measured global impact: the use of TZP and CTX/CRO predicted 3GC resistance and the use of TZP, IMP/MEM and, to a much lesser extent, CTZ/FEP predicted carbapenem resistance (Figure 3). The strikingly positive effect of TZP on both 3GCR and CR infections deserves further attention. Based on its in vitro efficacy against extended-spectrum beta-lactamase (ESBL) -producing enterobacteria, TZP has been repeatedly considered as an alternative drug of choice in carbapenem-sparing strategies^40,41^. The strategy of replacing carbapenems with TZP assumes that: (1), TZP treatment is as effective as carbapenems on TZP-susceptible pathogens; and (2) AMR is less selected under TZP pressure than under carbapenem pressure, as reflected by a recent consensus-based ranking of beta-lactams for deescalation therapy^16^. Yet, in several recent reports including a multicenter randomized clinical trial, TZP treatment of sepsis with ESBL-producing enterobacteria was associated with poorer outcome compared with carbapenem treatment^42–44^. Additionally, studies of the respective associations of TZP and carbapenem use with AMR yielded conflicting results. At the population level, the incidence of CR enterobacteria was negatively associated with TZP use in a 5-year, single-hospital trend analysis study^45^. At the patient level, however, TZP exposure was associated with CRPA in a meta-analysis^46^ and with CR Gramnegative bacilli in a single-hospital prospective cohort study^47^. Our finding that TZP use correlates with both 3GCR and CR infections, more strongly so than carbapenems and CTX/CRO, challenges the rationale of recommending TZP over other drugs for ecological reasons. Noteworthy, the link of TZP use with global 3GCR and CR resistance resulted from the accumulation of small, positive associations with most 3GCR and CR variants (Figure 2). Because of this diffuse effect, links between TZP use and 3GCR or CR infection incidence might go undetected in studies focusing on individual pathogen variants. It should be noted that our observation that TZP use selects for 3GCR and CR variants does not imply selection for acquired resistance through any specific mechanism such as carbapenemase production.

Our study has several limitations. Our inferences focused on infection incidence at the level of the ward and, as such, should not be readily translated to individual patients^19^. Because we modeled infection incidence, our analyses did not consider the contribution of asymptomatic carriers to AMR. We did not consider the movements of healthcare workers in our estimation of connectivity and therefore we could not determine their relative impact on AMR compared to patient transfers. We also modeled our network as a closed system, ignoring the effect of patient admissions from the community. The small sample sizes of VREF and CRAB limited our ability to draw robust inferences regarding these variants of utmost importance in other settings^48,49^. Finally, our findings reflect the AMR ecology of a Western European area, with generally lower prevalences of carbapenemase-producing pathogens and VREF than in other regions of the world.

To conclude, the metapopulation modeling of ESKAPE_2_ pathogens in hospital wards shows that connectivity has a measurable, variant-specific impact on infection incidence, which supports the need to tailor strategies against AMR depending on the targeted pathogen. Along with novel hospital-level insights into the driving forces of AMR, our work illustrates the application of the methodological framework of metapopulation ecology to the problem of hospital AMR. Applying this framework to other healthcare settings could help inform the local and regional antibiotic stewardship and infection control strategies.

## Methods

### Data collection and compilation

We obtained data on infection incidence from the information system of the Institut des Agents Infectieux, the clinical microbiology laboratory of the Hospices Civils de Lyon, a group of university hospitals serving the Greater Lyon urban area (~1.4 million inhabitants) of France. For each ward from October 1^st^, 2016 to September 30^th^, 2017, we extracted the number of deduplicated clinical samples (1 per patient, selected on first occurrence, after exclusion of screening samples) positive for at least one of the ESKAPE_2_ species (as determined using Vitek MS MALDI-ToF identification, bioMérieux), falling into one of the resistance variant categories defined in Table 1. Resistance was based on available results for susceptibility to, where applicable, ceftriaxone, cefotaxime, ceftazidime, cefepime, imipenem, meropenem, oxacillin and vancomycin. Antibiotic use in defined daily doses (ddd) of all systemic antibacterial drugs (ATC classification term J01), as well as of specific (groups of) molecules defined in Table 2, were extracted from the pharmacy department information system. For each pair of wards, the number of patient transfers was extracted from the hospital information system along with, for each ward, the number of beds, the type of medical activity and the number of patient admissions. Because of the aggregated nature of the data, informed consent was not sought, in accordance with French regulations. Our main response variable was the number of patients per ward infected with each pathogen variant, expressed as incidence over 1y.

### Sampling bias control

To correct for bias due to varying microbiological sampling frequencies and locations across wards, we computed for each pathogen variant the expected incidence explained by sampling alone, under the assumption of constant pathogen prevalence across wards. Sampling locations were assigned to 6 location groups, namely, skin and soft tissues, respiratory tract, urinary tract, digestive tract, vascular access devices, and sterile tissues (such as CSF and peripheral blood cultures). For each location group, the probability P(Variant | Location) that a sample is positive for a given pathogen variant was aggregated for all wards in the network. For each ward and location, we computed the average number of samples per sampled patient N(Location | Ward). For each patient in a ward, the probability of being tested positive at least once for the variant of interest was the complement of the probability that all samples from all locations remained negative, P(Variant | Ward)=1−∏_*l* ∈ Locations_[1 − P(Variant | *l*)]^N(*l* | Ward)^. Finally, the expected incidence of infections in a ward was computed as the patient-level probability of being tested positive at least once, times the number of tested patients N, written Incidence(Variant | Ward) = *N* × P(Variant | Ward).

Clearly, variations of the incidence control value between wards depend only on the number and locations of microbiological samples taken, thus reflecting the incidence and types of bacterial infections at ward level but not variations of pathogen community structure. This value, in turn, is expected to correlate with both the amount of antibiotics used and the probability of detecting patients, leading to spurious correlation between incidence and antibiotic use in unadjusted models. Indeed, the incidence control exhibited strong correlation with both the observed cumulative incidence of all bacteria (R^2^ = 0.96, p=10^−243^, Supplementary Figure 1) and the observed total antibiotic use (R^2^ = 0.34, p = 10^−33^, Supplementary Figure 2). These correlations remained significant for most pathogen variants and specific antibiotics. The incidence control was added as a covariate in all models predicting infection incidence. Such adjusted models, thus, predicted the incidence of infections in excess of what would be expected based on variations in sampling intensity alone.

### Connectivity and other ward characteristics

Habitat quality in wards was described using explanatory variables adapted from Hanski’s metapopulation models^22,23^, namely patch size and connectivity, along with additional variables capturing patient fragility and antibiotic selection pressure. We considered each ward within the hospital system as a distinct habitat patch, *i*. We used the number of beds as a measure of “patch area,” *A*_*i*_. The connectivity between wards was implemented as a proxy to the true but unobservable number of introductions of each variant in each ward during the study period^50^. To estimate this quantity, we measured directional, partial connectivity *S*_*j,i*_ from ward *j* to ward *i* as the number of patients transferred from *j* to *i*, times the observed probability that each patient tested positive for the variant. Partial connectivity, thus, was the expected number of positive patients transferred from *j* to *i*. Finally, connectivity for ward *i* was the sum of all directional connectivities, ***S***_*i*_=∑_*j*_***S***_*j,i*_.

Along with size and connectivity, wards were characterized by patient fragility and antibiotic consumption. Fragility was coded on a 3-level ordinal scale. Lower values were assigned to wards with more robust patients and higher values to wards with more fragile patients. General wards were coded as 0, intermediate (progressive) care units as 1, and intensive care and blood cancer units as 2. Antibiotic use was normalized by dividing with the number of beds in each ward and expressed in ddd/bed/y.

### Statistical analysis

The statistical unit was the individual ward (n=357) in all analyses. We used the asymptotic Simpson index^51^, also known as the Hunter-Gaston index^52^, to determine the probability that two random isolates of a given variant were isolated in the same ward or that two random doses of a given antibiotic were delivered in the same ward. The index is defined as [∑***n*_*i*_** (***n***_*i*_−1)]/[***N*** (***N***−1)] where ***n***_*i*_ is the number of isolates (or antibiotic doses) isolated (or delivered) in ward *i* and ***N***_*i*_ =∑***n***_*i*_ is the total number of isolates (or antibiotic doses). While typical applications of the Simpson index in ecology examine the distribution of sampled taxa relative to a sampling location, we examine the distribution of sampling locations relative to the taxa. Hence, we used the term ‘concentration index’ to avoid confusion with a diversity measure. The *iNext* R package was used to determine bootstrap-based 95% confidence intervals^53^.

Models of infection incidence were constructed using Poisson regression. All continuous explanatory variables including the incidence control, ward size, connectivity and antibiotic use were log_2_ transformed before further analyses. To avoid negative infinity values from this transformation, all zeroes were first converted to half the minimum non-zero value. This transformation was associated with better model fit (using the model structure of Figure 1), in terms of Akaike information criterion, compared with: (1) replacing zeroes with the minimum non-zero value before taking logs; (2) adding 1 to zero values before taking logs; or (3) avoiding log transformation. The log_2_ transformed data were used for all subsequent analyses. Analyses used R software version 3.6.0.

### Possibly causal associations between antibiotic use and resistance

We examined criteria to identify *a priori* possibly causal associations between antibiotic use and resistance. The criteria were based on medical and biological considerations, namely, that antibiotics inactive against a variant are unlikely to be prescribed in response to this variant’s prevalence; and that antibiotics are most likely to select for a variant when resistance provides a specific advantage, hence, when the variant is not resistant to more potent antibiotics (e.g, CTX/CRO is more likely to select for PA than for CRPA in which carbapenem resistance provides no additional benefit under CTX/CRO pressure). This rationale led to the following criteria: (1) the variant is always resistant to the antibiotics of interest; (2) the variant is not resistant to antibiotics more potent (in terms of spectrum or efficacy) than the antibiotics of interest; and (3) the antibiotics of interest can be plausibly used against the variant in empirical therapy. A total of 17 associations fulfilled the criteria for possible causality: IPM/MEM with CREC, CRKP, CREB, CRPA and CRAB; 1GC/2GC with MRSA; CTX/CRO with 3GCREC, 3GCRKP, 3GCREB, PA, AB and EF; CTZ/FEP with EF; VAN/TEC with VREF; OXA with MRSA; and AMC with EB and MRSA.

### Pooled analysis of CTX/CRO- and IPM/MEM-resistant variants

To model the cumulative incidences of 3GCR and CR infections, variants were pooled into resistance categories. When resistance to CTX/CRO or IPM/MEM was not determined by design (such as 3GC resistance in 3GCREC) or by intrinsic resistance (such as 3GC resistance in *E. faecium*), variants were classified as resistant when the proportion of resistance in our setting was above 80%. The rationale for this choice was that, contrary to the criteria for possible causality whose stringency was desirable to avoid ambiguity, excluding variants that are mostly resistant to an antibiotic group would bias pooled analyses. Applying the 80% threshold for the proportion of resistance led to classifying CRKP and CREB as 3GCR (91% and 93% 3GC resistance, respectively) but not CREC (61% 3GC resistance); and EF and VREF as CR (84% and 100% carbapenem resistance, respectively, inferred from ampicillin resistance^54^). Overall, the 3GCR category included 3GCREC, 3GCRKP, CRKP, 3GCREB, CREB, PA, CRPA, AB, CRAB, EF, VREF and MRSA; and the CR category included CREC, CRKP, CREB, CRPA, CRAB, EF, VREF and MRSA.

## Data Availability

All data that support the findings of this study are available from the corresponding author upon reasonable request.

## Code Availability

All code that support the findings of this study are available from the corresponding author upon reasonable request.

## Acknowledgements

The authors thank A. Friggeri, A. Lepape, F. Wallet and members of the LabEx EcoFect for fruitful discussion.

## Author Contributions

Conceived the study: JTS JPR JPF. Collected data: JPR GLe AFMD JFS PG AL MM OD SCT. Interpreted the data: JPR JTS GLe GLi CC FL FV. Analyzed the data:JTS JPR MM. Wrote the manuscript: JTS JPR. Revised the manuscript: GLe GLi CC FV BL JPF. Approved the final manuscript: all authors.

## Funding

This research received funding from the FINOVI Foundation (Grant R18037CC to JPR). The funder had no role in study design, data collection and analysis, manuscript preparation or decision to publish.

## Competing interests

The authors declare no competing interests.

## References

1. Laxminarayan, R. et al. Antibiotic resistance-the need for global solutions. Lancet Infect Dis 13, 1057–1098 (2013).

2. Cassini, A. et al. Attributable deaths and disability-adjusted life-years caused by infections with antibiotic-resistant bacteria in the EU and the European Economic Area in 2015: a population-level modelling analysis. Lancet Infect Dis 19, 56–66 (2019).

3. Johnson, T. A. et al. Clusters of Antibiotic Resistance Genes Enriched Together Stay Together in Swine Agriculture. MBio 7, e02214–02215 (2016).

4. Venter, H., Henningsen, M. L. & Begg, S. L. Antimicrobial resistance in healthcare, agriculture and the environment: the biochemistry behind the headlines. Essays Biochem. 61, 1–10 (2017).

5. Lübbert, C. et al. Environmental pollution with antimicrobial agents from bulk drug manufacturing industries in Hyderabad, South India, is associated with dissemination of extended-spectrum betalactamase and carbapenemase-producing pathogens. Infection 45, 479–491 (2017).

6. Chatterjee, A. et al. Quantifying drivers of antibiotic resistance in humans: a systematic review. Lancet Infect Dis (2018). doi:10.1016/S1473-3099(18)30296-2

7. David, S. et al. Epidemic of carbapenem-resistant Klebsiella pneumoniae in Europe is driven by nosocomial spread. Nat Microbiol (2019). doi:10.1038/s41564-019-0492-8

8. Safdar, N. & Maki, D. G. The commonality of risk factors for nosocomial colonization and infection with antimicrobial-resistant Staphylococcus aureus, enterococcus, gram-negative bacilli, Clostridium difficile, and Candida. Ann. Intern. Med. 136, 834–844 (2002).

9. Snitkin, E. S. et al. Tracking a hospital outbreak of carbapenem-resistant Klebsiella pneumoniae with whole-genome sequencing. Sci Transl Med 4, 148ra116 (2012).

10. Manning, M. L. et al. Antimicrobial stewardship and infection prevention-leveraging the synergy: A position paper update. Am J Infect Control 46, 364–368 (2018).

11. Goff, D. A. et al. A global call from five countries to collaborate in antibiotic stewardship: united we succeed, divided we might fail. Lancet Infect Dis 17, e56–e63 (2017).

12. Lemmen, S. W. & Lewalter, K. Antibiotic stewardship and horizontal infection control are more effective than screening, isolation and eradication. Infection 46, 581–590 (2018).

13. Chastain, D. B., White, B. P., Cretella, D. A. & Bland, C. M. Is It Time to Rethink the Notion of Carbapenem-Sparing Therapy Against Extended-Spectrum β-Lactamase-Producing Enterobacteriaceae Bloodstream Infections? A Critical Review. Ann Pharmacother 52, 484–492 (2018).

14. Robotham, J. V. et al. Cost-effectiveness of national mandatory screening of all admissions to English National Health Service hospitals for meticillin-resistant Staphylococcus aureus: a mathematical modelling study. Lancet Infect Dis 16, 348–356 (2016).

15. Kardaś-Słoma, L. et al. Universal or targeted approach to prevent the transmission of extended-spectrum beta-lactamase-producing Enterobacteriaceae in intensive care units: a cost-effectiveness analysis. BMJ Open 7, e017402 (2017).

16. Weiss, E. et al. Elaboration of a consensual definition of deescalation allowing a ranking of β-lactams. Clin. Microbiol. Infect. 21, 649.e1–10 (2015).

17. Acar, J. Broad- and narrow-spectrum antibiotics: an unhelpful categorization. Clin. Microbiol. Infect. 3, 395–396 (1997).

18. Huttner, B., Pulcini, C. & Schouten, J. De-constructing de-escalation. Clin. Microbiol. Infect. 22, 958–959 (2016).

19. Schechner, V., Temkin, E., Harbarth, S., Carmeli, Y. & Schwaber, M. J. Epidemiological interpretation of studies examining the effect of antibiotic usage on resistance. Clin. Microbiol. Rev. 26, 289–307 (2013).

21. Burton, D. C., Edwards, J. R., Horan, T. C., Jernigan, J. A. & Fridkin, S. K. Methicillin-resistant Staphylococcus aureus central line-associated bloodstream infections in US intensive care units, 1997-2007. JAMA 301, 727–736 (2009).

21. Levins, R. Some Demographic and Genetic Consequences of Environmental Heterogeneity for Biological Control. Bull Entomol Soc Am 15, 237–240 (1969).

22. Hanski, I. A Practical Model of Metapopulation Dynamics. Journal of Animal Ecology 63, 151–162 (1994).

23. Hanski, I. Metapopulation dynamics. Nature 396, 41 (1998).

24. Macpherson, J. L. & Bright, P. W. Metapopulation dynamics and a landscape approach to conservation of lowland water voles (Arvicola amphibius). Landscape Ecology 26, 1395–1404 (2011).

25. Heard, G. W. et al. Refugia and connectivity sustain amphibian metapopulations afflicted by disease. Ecology Letters 18, 853–863 (2015).

26. Dolrenry, S., Stenglein, J., Hazzah, L., Lutz, R. S. & Frank, L. A Meta-population Approach to African Lion (Panthera leo) Conservation. PLoS One 9, (2014).

27. Spagnolo, F., Cristofari, P., Tatonetti, N. P., Ginzburg, L. R. & Dykhuizen, D. E. Pathogen population structure can explain hospital outbreaks. ISME J 12, 2835–2843 (2018).

28. Vilches, T. N. et al. The role of intra and inter-hospital patient transfer in the dissemination of heathcare-associated multidrug-resistant pathogens. Epidemics (2018). doi:10.1016/j.epidem.2018.11.001

29. Hanski, I. & Gilpin, M. Metapopulation dynamics: Brief history and conceptual domain. Biological Journal of the Linnean Society 42, 3–16 (1991).

30. Bell, B. G., Schellevis, F., Stobberingh, E., Goossens, H. & Pringle, M. A systematic review and meta-analysis of the effects of antibiotic consumption on antibiotic resistance. BMC Infectious Diseases 14, 13 (2014).

31. Patrick, D. M. & Hutchinson, J. Antibiotic use and population ecology: How you can reduce your “resistance footprint”. CMAJ 180, 416–421 (2009).

32. Lipsitch, M. & Samore, M. H. Antimicrobial use and antimicrobial resistance: A population perspective. Emerg Infect Dis 8, 347–354 (2002).

33. Donker, T., Wallinga, J. & Grundmann, H. Patient Referral Patterns and the Spread of Hospital-Acquired Infections through National Health Care Networks. PLOS Computational Biology 6, e1000715 (2010).

34. Donker, T. et al. Measuring distance through dense weighted networks: The case of hospital-associated pathogens. PLOS Computational Biology 13, e1005622 (2017).

35. Vilches, T. N. et al. The role of intra and inter-hospital patient transfer in the dissemination of heathcare-associated multidrug-resistant pathogens. Epidemics 26, 104–115 (2019).

36. Seigal, A., Mira, P., Sturmfels, B. & Barlow, M. Does Antibiotic Resistance Evolve in Hospitals? Bull. Math. Biol. 79, 191–208 (2017).

37. Cusini, A. et al. Intra-hospital differences in antibiotic use correlate with antimicrobial resistance rate in Escherichia coli and Klebsiella pneumoniae: a retrospective observational study. Antimicrob Resist Infect Control 7, 89 (2018).

38. Willemsen, I., Bogaers-Hofman, D., Winters, M. & Kluytmans, J. Correlation between antibiotic use and resistance in a hospital: temporary and ward-specific observations. Infection 37, 432–437 (2009).

39. Huang, T.-D. et al. Temocillin and piperacillin/tazobactam resistance by disc diffusion as antimicrobial surrogate markers for the detection of carbapenemase-producing Enterobacteriaceae in geographical areas with a high prevalence of OXA-48 producers. J. Antimicrob. Chemother. 69, 445–450 (2014).

40. Harris, P. N. A., Tambyah, P. A. & Paterson, D. L. β-lactam and β-lactamase inhibitor combinations in the treatment of extended-spectrum β-lactamase producing Enterobacteriaceae: time for a re-appraisal in the era of few antibiotic options? The Lancet Infectious Diseases 15, 475–485 (2015).

41. Peterson, L. R. Antibiotic policy and prescribing strategies for therapy of extended-spectrum β-lactamase-producing Enterobacteriaceae: the role of piperacillin–tazobactam. Clinical Microbiology and Infection 14, 181–184 (2008).

42. Ofer-Friedman, H. et al. Carbapenems Versus Piperacillin-Tazobactam for Bloodstream Infections of Nonurinary Source Caused by Extended-Spectrum Beta-Lactamase-Producing Enterobacteriaceae. Infect Control Hosp Epidemiol 36, 981–985 (2015).

43. Tamma, P. D. et al. Carbapenem therapy is associated with improved survival compared with piperacillin-tazobactam for patients with extended-spectrum β-lactamase bacteremia. Clin. Infect. Dis. 60, 1319–1325 (2015).

44. Harris, P. N. A. et al. Effect of Piperacillin-Tazobactam vs Meropenem on 30-Day Mortality for Patients With E coli or Klebsiella pneumoniae Bloodstream Infection and Ceftriaxone Resistance: A Randomized Clinical Trial. JAMA 320, 984–994 (2018).

45. McLaughlin, M. et al. Correlations of antibiotic use and carbapenem resistance in enterobacteriaceae. Antimicrob. Agents Chemother. 57, 5131–5133 (2013).

46. Raman, G., Avendano, E. E., Chan, J., Merchant, S. & Puzniak, L. Risk factors for hospitalized patients with resistant or multidrug-resistant Pseudomonas aeruginosa infections: a systematic review and meta-analysis. Antimicrob Resist Infect Control 7, 79 (2018).

47. Marchenay, P. et al. Acquisition of carbapenem-resistant Gramnegative bacilli in intensive care unit: predictors and molecular epidemiology. Med Mal Infect 45, 34–40 (2015).

48. Arias, C. A. & Murray, B. E. The rise of the Enterococcus: beyond vancomycin resistance. Nat. Rev. Microbiol. 10, 266–278 (2012).

49. Hsu, L.-Y. et al. Carbapenem-Resistant Acinetobacter baumannii and Enterobacteriaceae in South and Southeast Asia. Clin. Microbiol. Rev. 30, 1–22 (2017).

50. Donker, T. et al. Measuring distance through dense weighted networks: The case of hospital-associated pathogens. PLOS Computational Biology 13, e1005622 (2017).

51. Simpson, E. H. Measurement of Diversity. Nature 163, 688 (1949).

52. Hunter, P. R. & Gaston, M. A. Numerical index of the discriminatory ability of typing systems: an application of Simpson’s index of diversity. J. Clin. Microbiol. 26, 2465–2466 (1988).

53. Hsieh, T. C., Ma, K. H. & Chao, A. iNEXT: an R package for rarefaction and extrapolation of species diversity (Hill numbers). Methods in Ecology and Evolution 7, 1451–1456 (2016).

54. Weinstein, M. P. Comparative evaluation of penicillin, ampicillin, and imipenem MICs and susceptibility breakpoints for vancomycin-susceptible and vancomycin-resistant Enterococcus faecalis and Enterococcus faecium. J. Clin. Microbiol. 39, 2729–2731 (2001).

