## Supplementary Information for "Metapopulation ecology links antibiotic resistance, consumption and patient transfers in a network of hospital wards"

**Supplementary Information File**

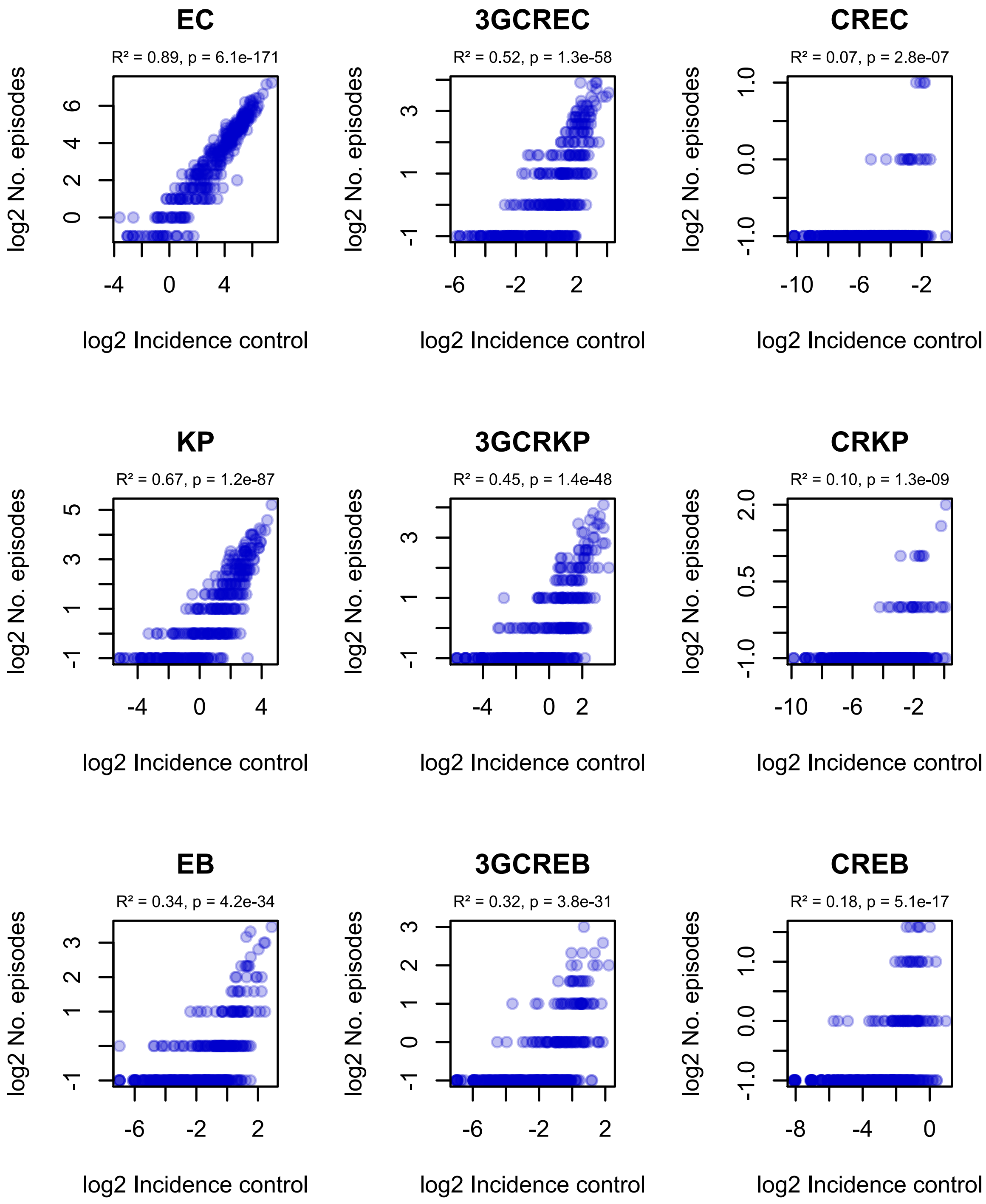


**Supplementary Figure 1. Correlation of ward-level incidence control values with infection incidence in ESKAPE$_2$ variants.** $R^2$ and p-values were obtained using simple linear regression on log2-transformed data. Figure continues on next page.

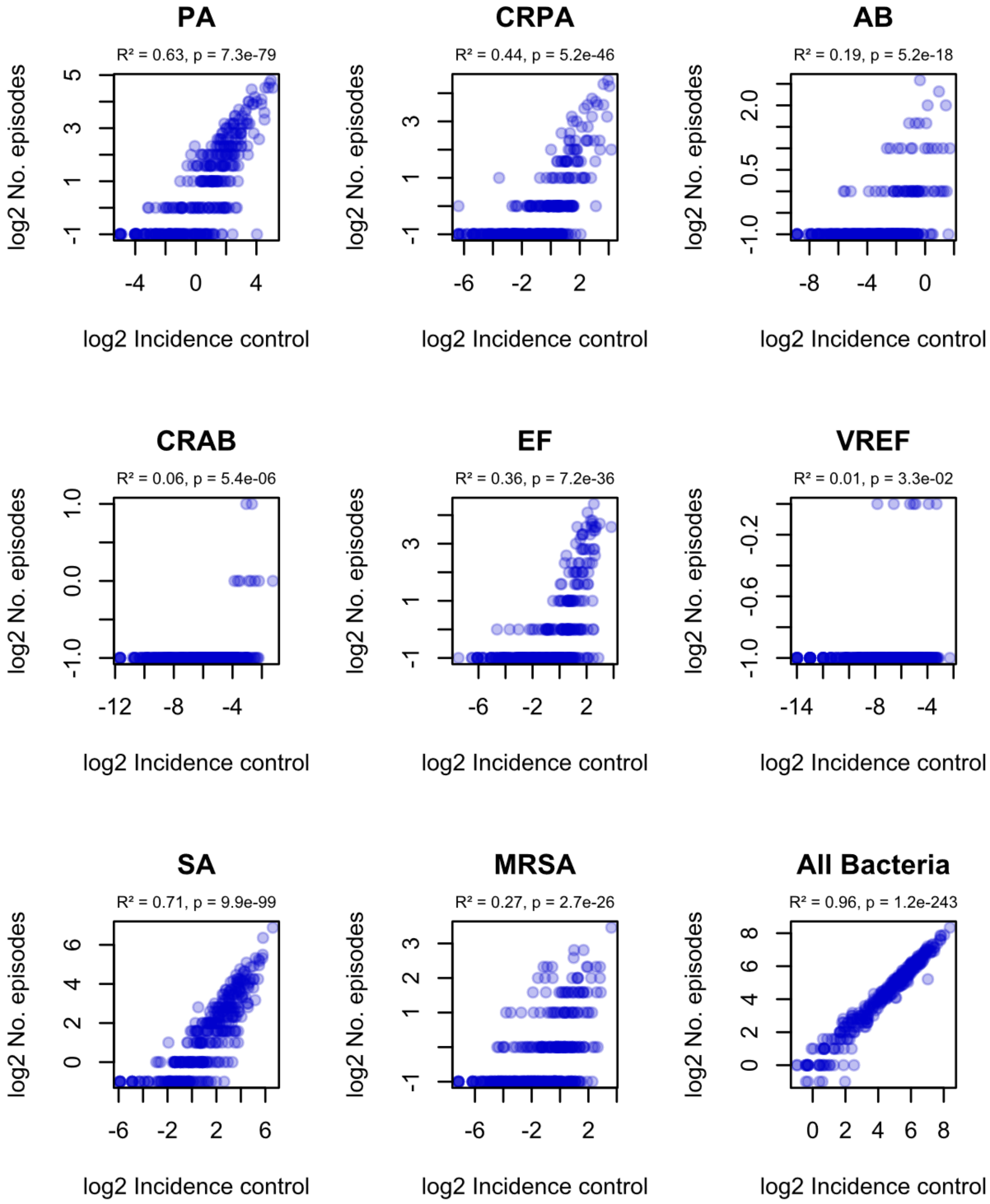


**Supplementary Figure 1 (continued). Correlation of ward-level incidence control values with infection incidence in $ESKAPE_2$ variants.** $R^2$ and p-values were obtained using simple linear regression on log2-transformed data.

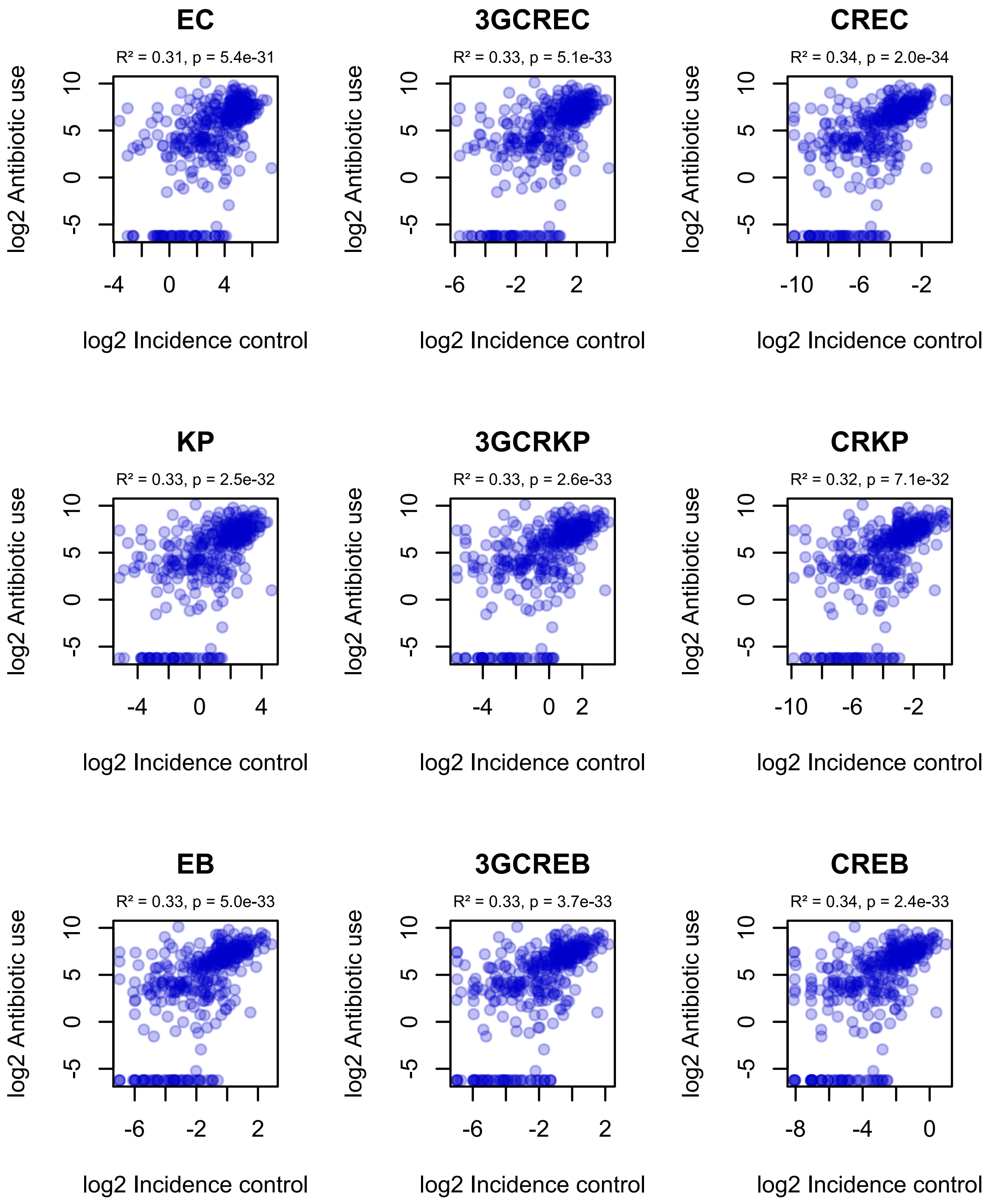


**Supplementary Figure 2. Correlation of incidence control values with observed ward-level antibiotic consumption.** $R^2$ and p-values were obtained using simple linear regression on log2-transformed data. Figure continues on next page.

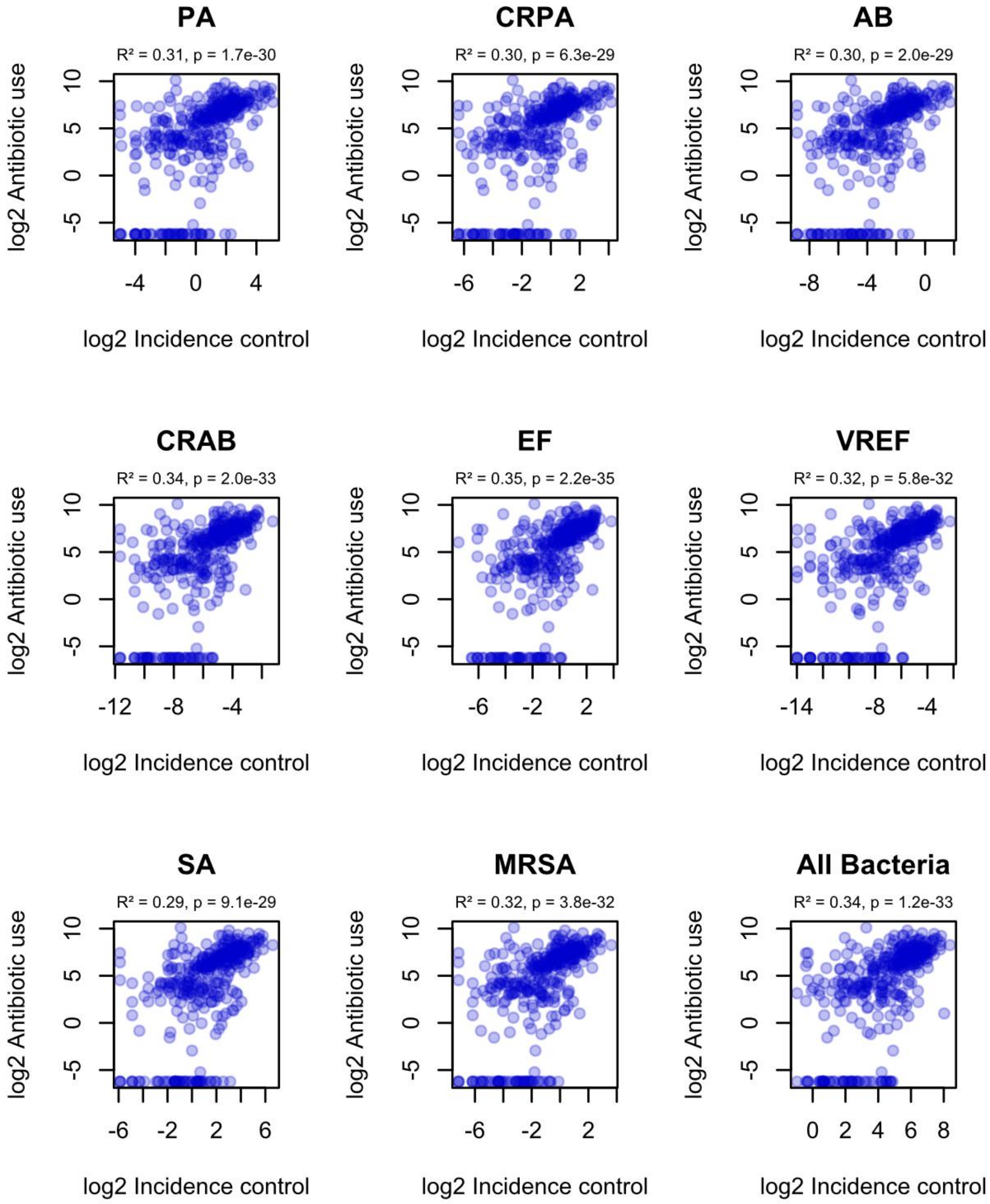


**Supplementary Figure 2 (continued). Correlation of incidence control values with observed ward-level antibiotic consumption.** $R^2$ and p-values were obtained using simple linear regression on log2-transformed data.

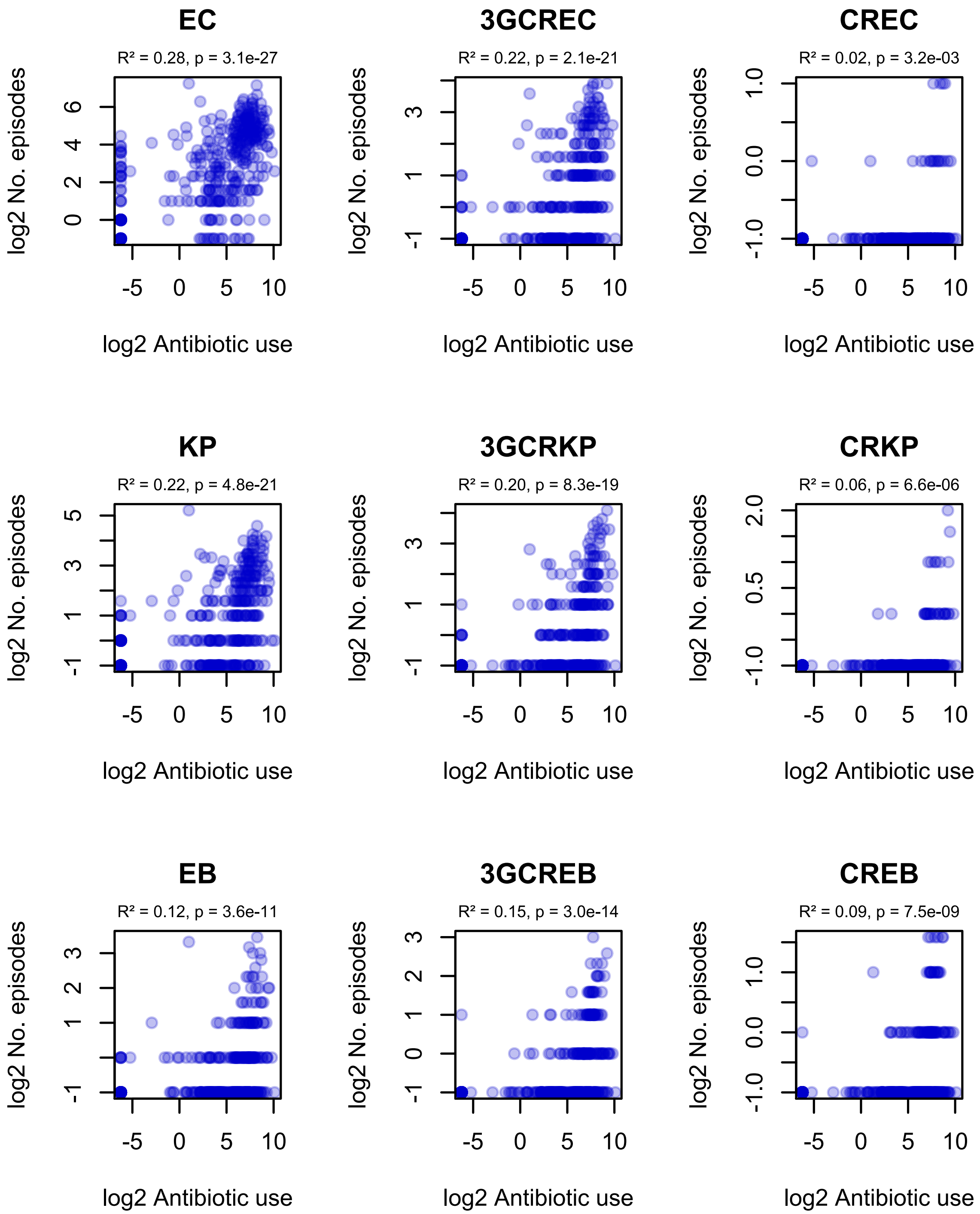


**Supplementary Figure 3. Correlation of ward-level antibiotic consumption and infection incidence in ESKAPE$_2$ variants.** $R^2$ and p-values were obtained using simple linear regression on log2-transformed data. Figure continues on next page.

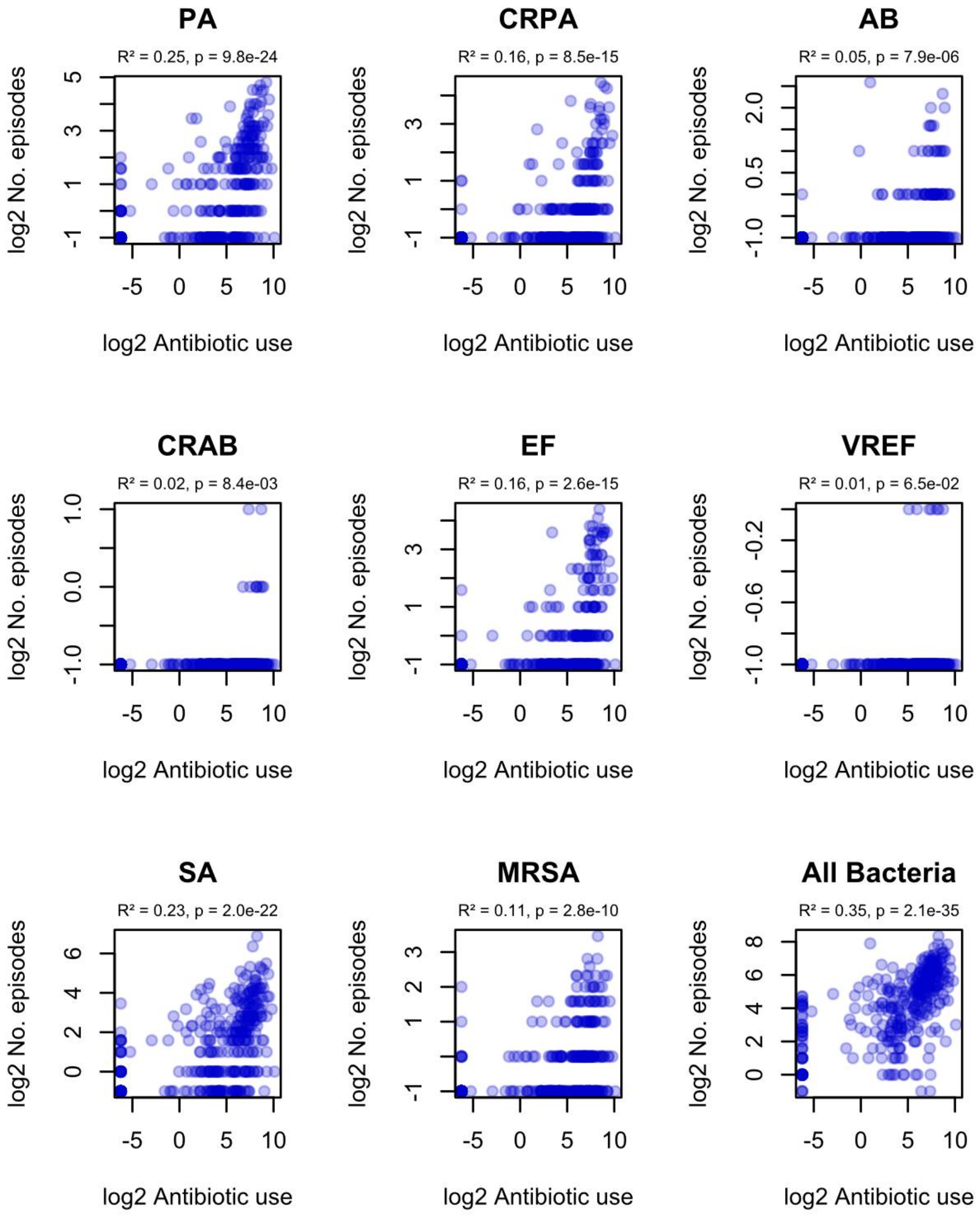


**Supplementary Figure 3 (continued). Correlation of ward-level antibiotic consumption and infection incidence in $ESKAPE_2$ variants.** $R^2$ and p-values were obtained using simple linear regression on log2-transformed data.

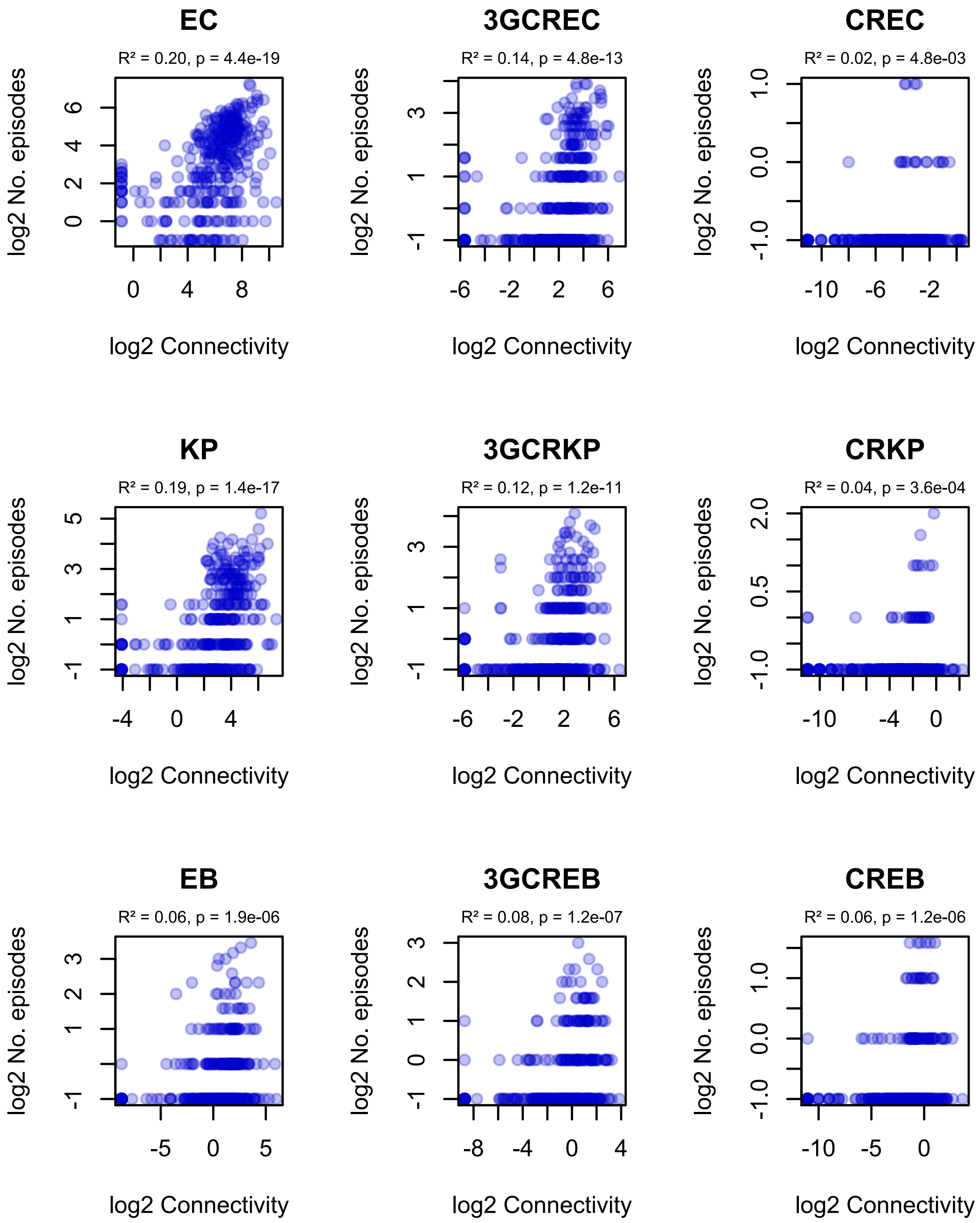


**Supplementary Figure 4. Correlation of ward-level connectivity and infection incidence in ESKAPE$_2$ variants.** $R^2$ and p-values were obtained using simple linear regression on log2-transformed data. Figure continues on next page.

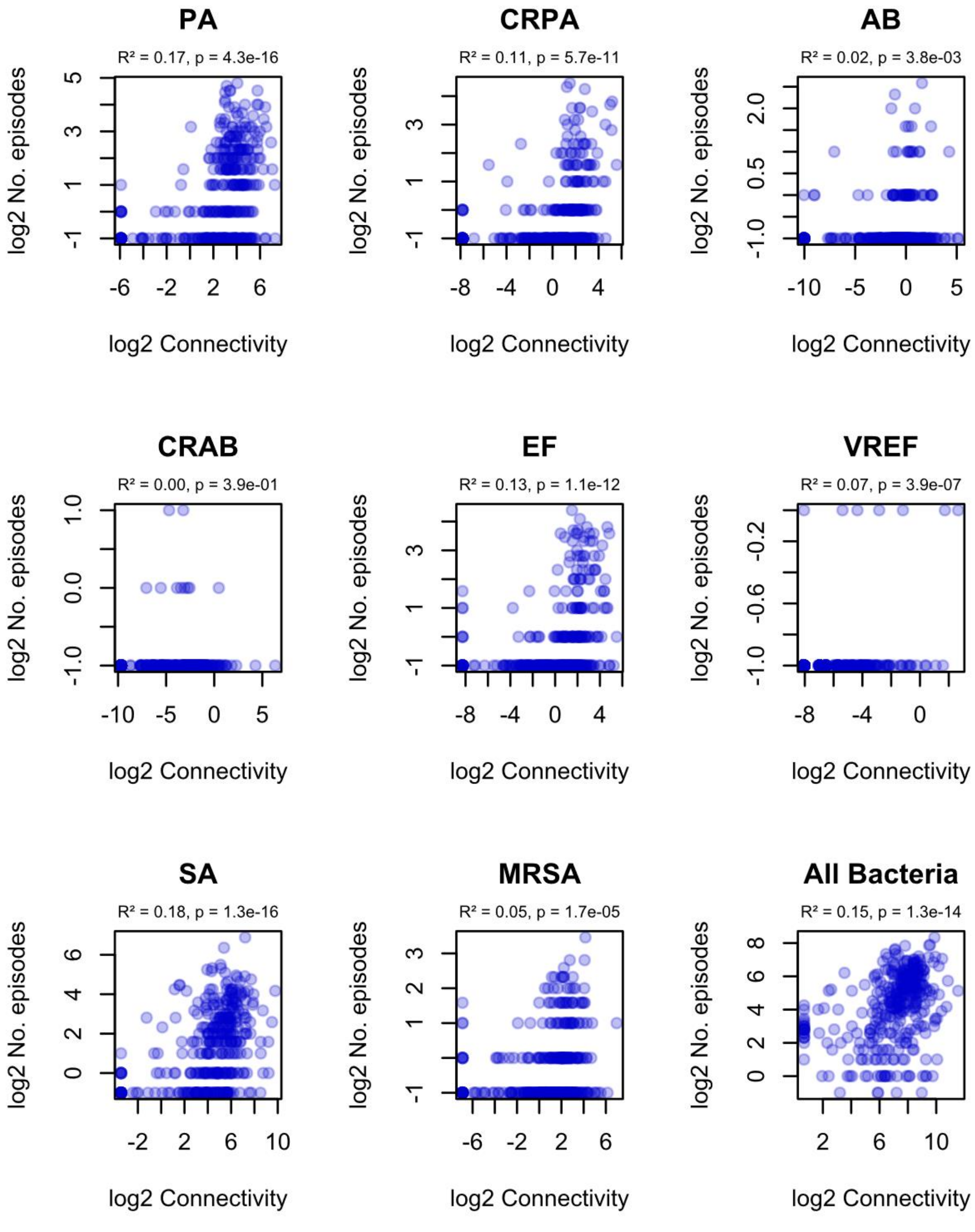


**Supplementary Figure 4 (continued). Correlation of ward-level connectivity and infection incidence in ESKAPE$_2$ variants.** $R^2$ and p-values were obtained using simple linear regression on log2-transformed data.
